# Hyperoxia Induced Alteration of Chromatin Structure in Bone Marrow Derived Primary Mesenchymal Stromal Cells

**DOI:** 10.1101/2023.12.29.573604

**Authors:** Lauren A. Monroe, Samantha Kaonis, Natalie Calahan, Neda Kabi, Soham Ghosh

**Author notes:** Corresponding author: Soham Ghosh.

## Abstract

Chromatin is a highly dynamic entity of the eukaryotic cell nucleus. New evidence is emerging in support of the notion that chromatin can locally and globally rearrange itself to adapt with the cellular microenvironmental changes. Such changes include oxidative stress such as supraphysiological oxygen level, found in hyperoxia. Although it is known that hyperoxia can result in DNA damage and alterations in cell function, it is not well understood how the chromatin architecture changes under such a condition and what the functional significance of such change entails. In this work we developed an imaging-based technique to visualize and characterize nanoscale chromatin remodeling under hyperoxia, created via hydrogen peroxide treatment. We found high spatiotemporal variability of remodeling in different chromatin domains such as the euchromatin, heterochromatin and interchromatin. Chromatin remodeling was hindered by the GSK126 mediated inhibition of methyltransferase EZH2, which regulates the chromatin compaction. Epigenetic modifications and DNA damage under hyperoxia was investigated, which was found affected by the pretreatment of GSK126. The developed techniques and findings inform us with new mechanistic insights of chromatin remodeling which might lead to new intervention strategies to target genotoxic hyper-oxidative stress, which is common in degenerative diseases and aging, and for cell therapy in regenerative medicine.

## INTRODUCTION

Cells accumulate DNA damage throughout their lifetime due to natural processes like cellular metabolism, where reactive oxygen species (ROS) are by-products. As organisms age, ROS levels may increase, leading to more DNA damage. Environmental stresses common in physiological systems also generate ROS. This hyper-oxidative condition, or hyperoxia, results from the generation and accumulation of ROS, damaging cellular macromolecules, including DNA, and causing cell apoptosis and senescence[1]. With aging and tissue degeneration, the hyper-oxidative stress intensifies[2–6]. Therefore, the combined effects of aging and environmental stress lead to DNA damage through increased hyper-oxidative stress. Cells demonstrate various responses upon incurring DNA damage that include the repairing the double stranded break. In epithelial and other cells, chromatin is shown to remodel to protect the nucleus from the adverse effects of DNA damage lesions [7–10]. Euchromatin, the relatively open region in the nucleus and heterochromatin, the denser regions in the nucleus were proposed to play a critical role in damage correction. DNA damage triggers the formation of a dense heterochromatin barrier, but the cause of such a response is unknown. Conflicting reports exist where some studies suggest that the dense heterochromatin creates a barrier around the lesions to prevent the damage spreading into the nucleus while others claim that damaged DNA lesion may be carried to an open euchromatin region for efficient damage correction because damage correction resources are abundant in the euchromatin region[10,11]. Therefore, it is not understood what the steps post exposure to hyper-oxidative stress are which facilitate such chromatin remodeling. Additionally, the role of specific epigenetic modifiers in chromatin remodeling in response to ROS is unknown. In this study, we aim to discover these underlying mechanisms using advanced microscopy and image analysis techniques using mesenchymal stem/ stromal cells (MSC) as the primary model.

Oxidative stress is particularly critical for stromal cells such as MSC. MSC are residents in many tissues and they are known to play a role in tissue regeneration and immunomodulation [4,12]. During their lifetime they are continuously exposed to environmental stresses including ROS. MSCs are activated under such physiological stress. Accordingly, low dose of biochemical and biophysical stress inducing conditions have been exploited to mobilize and activate MSC for enhanced regeneration *in vivo* [13–15]. Under these conditions, MSC show higher activation, differentiation and proliferation [16–24]. However, under prolonged and repeated stress inducing events, MSC can become senescent and lose their functionality. Therefore, maintenance of optimal oxidative tension status is critical for tissues and for MSC to survive and regenerate during its lifetime [25–28]. Optimal oxidative stress is equally important for MSC based therapy as well. For regenerative medicine applications, culture expanded MSC are injected inside the damaged tissue where they are exposed to high amounts of oxygen toxicity [29,30], associated with inflammatory conditions. Therefore, although it is evident that MSC and other cells are exposed to hyper oxidative stress in vivo and in vitro, it’s not clear what happens inside MSC or other cell types at the chromatin level after the exposure to hyper-oxidative stress due to technical limitations to investigate such events. Understanding such mechanisms might provide us with new mechanisms and intervention strategies to target genotoxic stress, which is common in degenerative diseases and aging, and for cell therapy in regenerative medicine.

The dynamic nature of the intranuclear space within cells plays a crucial role in gene expression regulation over time, with chromatin remodeling emerging as a key mechanism. This remodeling is facilitated by various epigenetic processes, such as through the Chromatin Remodeling Complex (CRC) and histone modifications, which locally condense or decondense chromatin. Although the detailed mechanism and functional significance of chromatin remodeling have recently come to light, a specific type of remodeling occurs at the multi-nucleosome level, where ATP-dependent chromatin remodelers play a critical role. The intricate details of chromatin remodeling, involving processes like nucleosome sliding, engagement, and ejection, are still being uncovered. Various classes of CRC, including SWI/SNF (<u>SWI</u>tch/ <u>S</u>ucrose <u>N</u>on-<u>F</u>ermentable), have been identified. The core protein in the SWI/SNF complex, ARID1A governs chromatin remodeling by nucleosome sliding and ejection. Genes are expressed by the combined effects of ARID1A and a histone methyltransferase enzyme EZH2[31]. EZH2 is responsible for the methylation activity of Polycomb Repressive Complex 2 (PRC2), especially the addition of methyl groups to histone H3 at lysine 27[32] thus determining the H3K27Me3 level. Consequently, pharmacological intervention of EZH2 offers an opportunity to explore the role of chromatin remodeling in cellular functions, especially during sudden perturbations that may necessitate chromatin compaction and remodeling.

In this study, we hypothesized that hyper oxidative stress in cells triggers chromatin remodeling by chromatin condensation. To test this hypothesis, we visualized and characterized the chromatin remodeling at high spatiotemporal resolution in cells undergoing hyper-oxidative stress by hydrogen peroxide using a newly developed imaging assay. Chromatin remodeling was impeded by the inhibition of methyltransferase EZH2, which regulates the chromatin compaction with the drug GSK126. The drug has a very high affinity and exhibits >150-fold selectivity for EZH2 over EZH1 and >1000-fold selectivity over twenty other human methyltransferases. After thorough characterization of how GSK126 affects the MSC phenotype and several epigenetic modifiers, the effect of GSK126 on chromatin remodeling under hyperoxia was investigated. Finally, DNA damage was visualized in nuclei under hyper-oxidative stress and by the inhibition of chromatin remodeling using GSK126. Human bone marrow derived primary MSC were used for the majority of this study but murine embryonic fibroblast cell line, NIH 3T3, was also used to show the generic observations at the chromatin level. The results are discussed to provide novel insights into the role of chromatin remodeling and epigenetic changes during oxygen toxicity, with possible implications in cellular physiology in health, disease and regenerative medicine.

## RESULTS

### GSK126 lowers the cell proliferation by affecting chromatin compaction but does not affect cell phenotype

Quantification of cell proliferation by imaging revealed a significant reduction in MSC proliferation (hBM-MSC: human bone marrow derived mesenchymal stromal cells) on the application of 20 µM GSK126, as shown in Figure 1A. Notably, there were no discernible visual alterations in cell or nuclear phenotype between the control and GSK126-treated groups. The proliferation rate decresaed by nearly 60% over a 3-day period with GSK126 treatment, as illustrated in Figure 1B and 1C. Despite this reduction, MSCs maintained their characteristic spindle-like shape and elliptical nucleus in both experimental conditions, as quantified using conventional geometric parameters (Figure 1D, E) [33]. Traditional MSC specific markers (CD73 and CD105) did not display lower expression in the GSK126 treated group at two concentrations (Figure 1F). We observed the same trends with the proliferation of NIH 3T3 cells (Fig. S1), confirming the similar effect of GSK126 on two types of cells. GSK126 inhibits EZH2 therefore hindering H3K27 methylation which is asssociated with condensed chromatin. Overall, during the mitosis process heterochromatin formation is required and is facilitated by EZH2. Therefore, the loss of cell proliferation by the application of GSK126 is expected. However, although previous studies indicated that EZH2-PRC2-H3K27Me3 axis governs the gene silencing after cell division[34], our data suggests that the mitosis process in itself is hindered by EZH2 inhibition.

**Figure 1.**
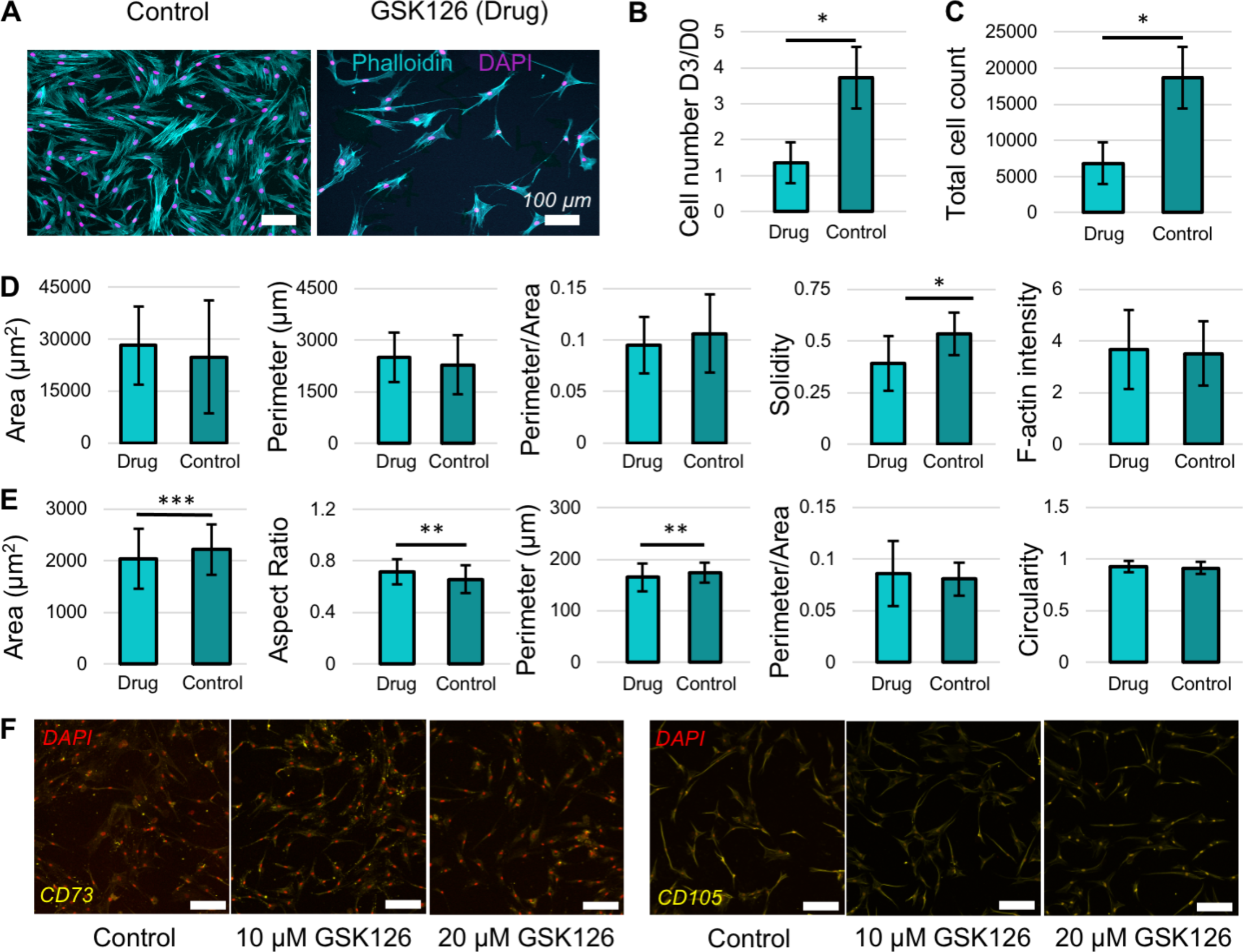
Effect of 20 µM GSK126 treatment on the proliferation and phenotype of human bone marrow derived primary MSC (hBM-MSC). **(A)** Representative images of Passage 4 MSC with phalloidin stained F-actin and DAPI stained nucleus on Day 3 after seeding. **(B)** Ratio of number of cells on Day 3 over Day 0. **(C)** Total number of cells on Day 3. **(D)** Different geometric shape parameters based upon cell phenotype. **(E)** Different geometric shape parameters based upon nuclear phenotype. Data based upon >4 technical replicates (>2000 cells per replicate) per group. *p <0.0001, **p <0.001, ***p <0.05. **(F)** Immunofluorescence imaging of two classic MSC markers CD73 and CD105 along with nuclear stain DAPI. Scale bar: 100 µm.

### H3K27Me3 level decreases, H3K4Me3 level remains the same, ARID1A level slightly increases by GSK126

GSK126, through its inhibition of EZH2, exerts an impact on chromatin remodeling and H3K27 methylation as known from previous studies[32]. Consequently, it is anticipated that this influence may extend to various epigenetic modifications such as H3K4Me3 and H3K9Me3. Previous studies suggest that in some cell types ARID1A and EZH2 affect gene expression in an antagonistic manner[31]. We investigated if EZH2 inhibition directly affects the ARID1A level in MSC. To explore this possibility, confocal microscopy was utilized to visualize H3K27Me3, H3K4Me3 and ARID1A levels, and the results are illustrated in Figure 2.

**Figure 2.**
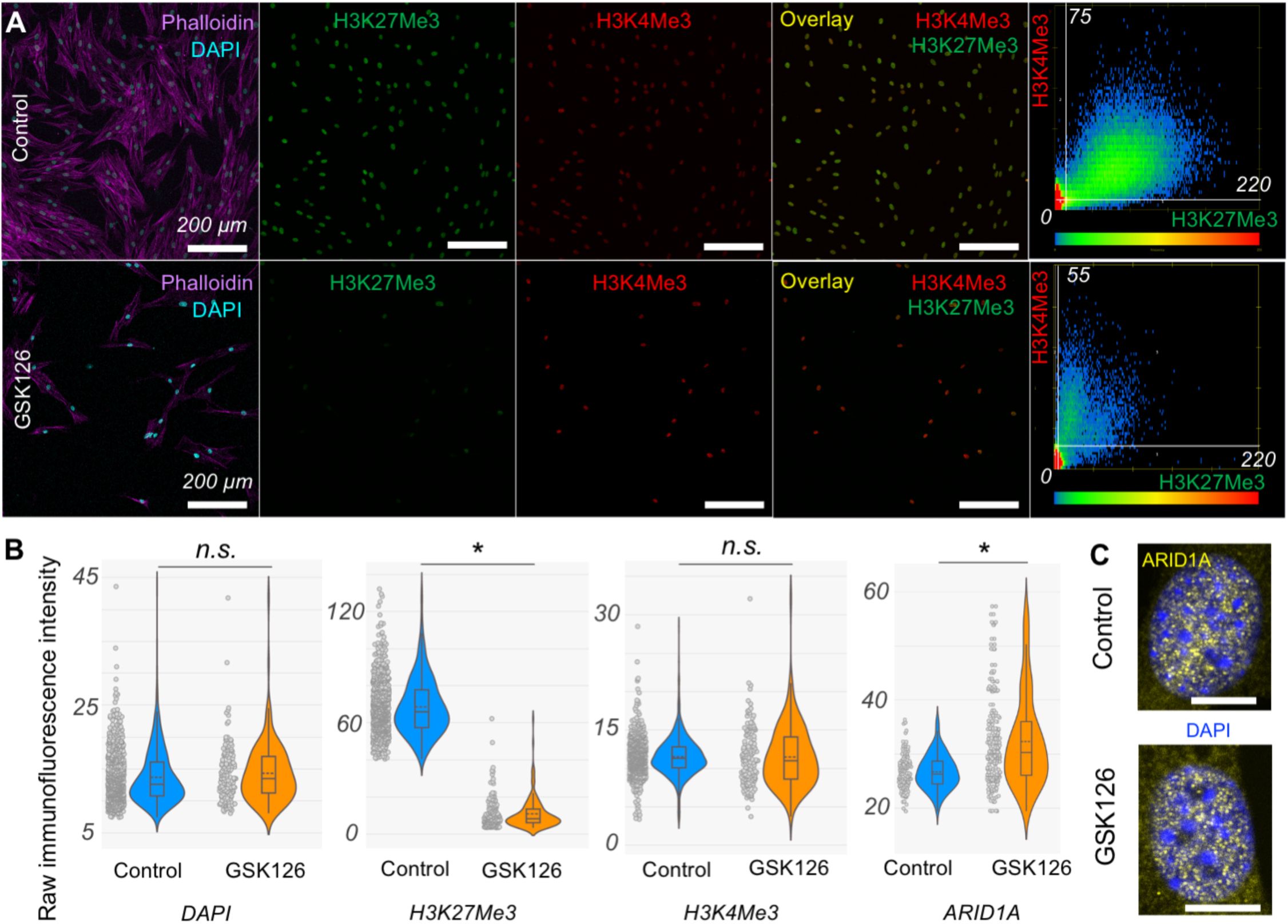
Effect of 20 µM GSK126 treatment on several epigenetic modifiers in hBM-MSC. **(A)** For both control and GSK126 treated groups, gene repression specific modifier H3K27Me3 and gene activation specific modifier H3K4Me3 are shown in unprocessed representative images to demonstrate their relative intensity levels. The scatterplot at the right panel shows the colocalization of H3K27Me3 and H3K4Me3 in the representative images. **(B)** Unprocessed image Intensity based quantification of the intensity of epigenetic modifiers. Data based upon 4 technical replicates (>30 cells per replicate for DAPI, H3K27Me3 and H3K4Me3 and >15 cells per replicate for ARID1A) per group. *p <0.001. All the intensity of the nuclear markers – H3K27Me3, H3K4Me3 and ARID1A are based upon the intensity of the stains in the nucleus, which was delineated from the DAPI channel. **(C)** High resolution confocal images of ARID1A and DAPI stained nuclei. Scale bar: 10 µm.

H3K4Me3, a specific epigenetic modifier associated with transcriptionally active regions, exhibited consistent intensity under GSK126 treatment. In contrast, H3K27Me3, the epigenetic modifier linked to repressed genes, displayed a nearly sixfold reduction in levels as reported in previous studies (Figure 2B) [31]. Furthermore, colocalization analysis using Manders’ Overlap Coefficient indicated a decrease in the simultaneous occurrence of H3K4Me3 and H3K27Me3 after a 3-day GSK126 treatment (Figure 2A). H3K27Me3 is a facultative repression marker, which marks ‘partially close’ chromatin and H3K4Me3 marks open chromatin. Therefore, this data suggests an overall open chromatin architecture upon EZH2 inhibition although the DAPI staining shows an overall similar intensity level (leftmost panel in Figure 2B). H3K9Me3 level change is shown and discussed later (Figure 6).

Interestingly, ARID1A intensity (Figure 2B) increased with GSK126 treatment, although the change was not as drastic as the decrease in H3K27Me3 level, still the distribution clearly indicates that a significant number of cells showed higher level of ARID1A in the nucleus. Surprisingly, higher resolution confocal microscopy images revealed that GSK126 did not affect the intranuclear spatial localization of ARID1A. Notably, ARID1A predominantly localized in euchromatin regions (Figure 2C) - the zones excluded by heavily DAPI-stained areas (heterochromatin) and areas excluded by DAPI-unstained regions, potentially populated by the nucleolus. This pattern remained consistent in both the control and GSK126-treated groups. Overall, this data suggests that although the nuclear ARID1A level slightly increases in the nucleus upon EZH2 inhibition, its distribution which might affect the chromatin remodeling does not change.

### Hydrogen peroxide causes nuclear compaction which is inhibited by GSK126 treatment

Hyperoxia, characterized by elevated levels of reactive oxygen species beyond physiological norms, is recognized for its detrimental effects on cellular integrity. In this study, we sought to examine the impact of hydrogen peroxide exposure on cell nuclei and chromatin through live microscopy of individual cell nuclei stained with NucBlue, a live cell specific formulation of Hoechst. As depicted in Figure 3, nuclei in the control group exhibited shrinkage following exposure to 500 µM H_2_O_2_ treatment, whereas nuclei in the GSK126 pre-treated group maintained a relatively constant area. The accompanying violin plots quantifies the compaction as the % decrease of nuclear area. The observed condensation in both cases persisted for approximately 75 to 80 minutes, after which nuclear size remained constant for another 30 minutes. We do not know how the nuclear area changes after this timepoint because live imaging was stopped after around 2 hours. Experiments with fibroblasts showed the same trend which further confirms such hyperoxia induced chromatin condensation event (Fig. S2). This data highlights the phenomenon of chromatin condensation in response to H_2_O_2_ exposure, underscoring the essential role of chromatin remodeling in this process. Furthermore, it was evident that chromatin compaction is a key requirement in this process and the disruption of chromatin remodeling by EZH2 inhibition can hinder the nuclear area shrinkage.

**Figure 3.**
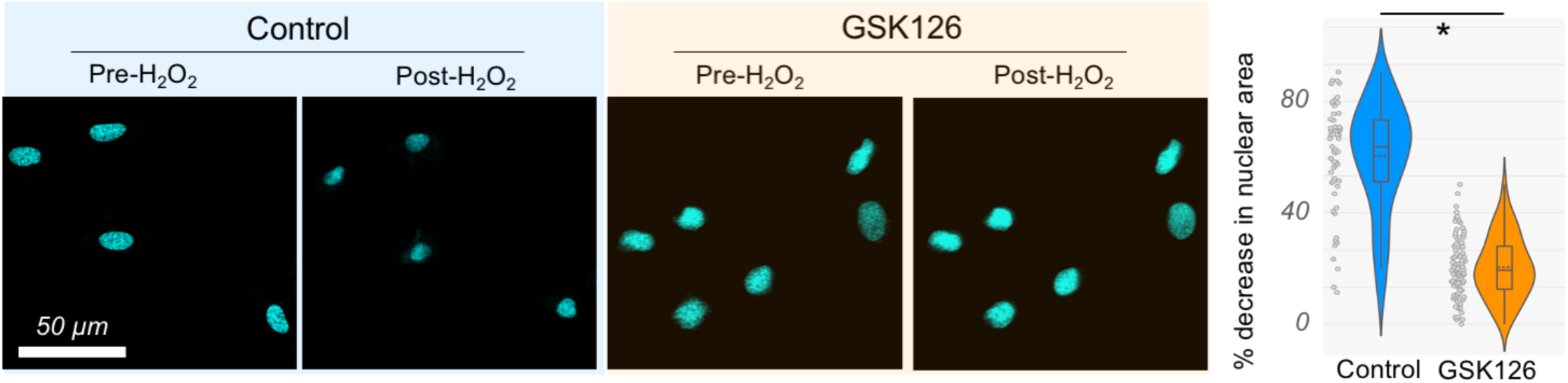
H_2_O_2_ (500 µM) driven nuclear shrinkage in hBM-MSC. Representative Images of Nucblue stained nuclei in live cells before and after the application of H_2_O_2_. Post-H_2_O_2_ corresponds to 75 min timepoint. Nucblue area decreases as a result of chromatin condensation in the control group. GSK126 treated groups show no such decrease in the chromatin condensation and nuclear area. Data based upon > 50 cells per group. *p <0.01.

### Chromatin remodeling is zone specific inside the nucleus

Utilizing high-resolution time-lapse imaging of individual nuclei stained with NucBlue during exposure to H_2_O_2_, active chromatin remodeling was observed, as shown in Figure 4A. For the time duration of imaging, we did not notice any significant amount of photobleaching because with the airyscan imaging modality, the laser power was kept extremely low < 0.2% and the exposure time was also very low without compromising the imaging quality. The colormap shows that although there was an overall shrinkage of the nucleus upon H_2_O_2_ application, the chromatin remodeling was spatially very heterogeneous. Notably, the remodeling was significantly halted by the pre-treatment with GSK126. Chromatin Remodeling Index (CRI) maps also illustrated the temporal variability in the remodeling process. To discern the chromatin zone within the nucleus undergoing the most pronounced remodeling, the intranuclear space was partitioned into three regions: euchromatin (open region), heterochromatin (closed region), and interchromatin (interface of heterochromatin and euchromatin) by modifying a technique we reported in a prior work[35].

**Figure 4.**
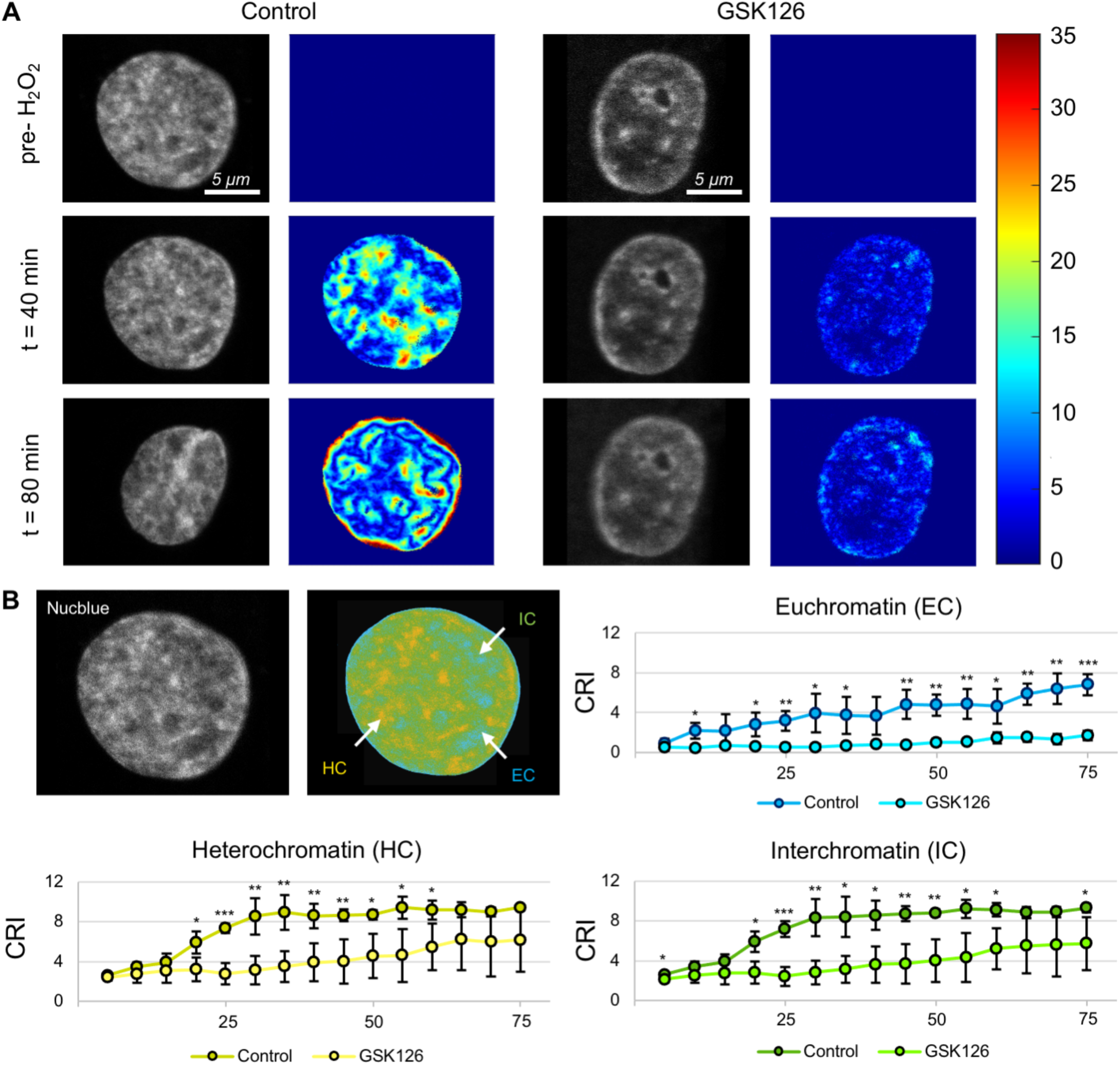
Chromatin remodeling in hBM-MSC under the exposure of 500 µM H_2_O_2_. **(A)** Time lapse image sequence at a few timepoints for representative nuclei without or with GSK126 treatment. Chromatin Remodeling Index map (displacement in pixels) is shown as colormap. 1 pixel = 21 nm. **(B)** Heterochromatin (HC), Euchromatin (EC) and Interchromatin (IC) domains shown for a representative nucleus. In the plots, X axis represent time (in minutes) and Y axis represents CRI in the specific domain. Data based on at least 5 nuclei per group. * p <0.05 **p <0.01, ***p <0.001.

The pixel resolution is around 20 nm and locally, the chromatin remodeling occurs at the similar scale which reveals a nanoscale chromatin reorganization or displacement although from the time lapse images it looks like a global process. Time-lapse average CRI values for each zone, as shown in Figure 4B, revealed that the euchromatin domain experienced the most substantial chromatin remodeling, particularly when compared to the GSK126 treated group. Additionally, chromatin remodeling in euchromatin persisted for an extended duration in comparison to the flatter time-lapse curves observed in the heterochromatin and interchromatin regions. This data suggests that chromatin compaction mostly occurs and sustains at the already open euchromatin regions. Additionally, the data suggests that GSK126 impedes the nanoscale chromatin remodeling process occurring during H_2_O_2_ exposure.

### H3K9Me3 level is differentially affected by GSK126 and hydrogen peroxide treatment

H3K9Me3 serves as an epigenetic modifier largely linked to constitutive heterochromatin, indicating its association with repressed regions that are not easily amenable to alteration. Upon visualizing and quantifying the intensity level of H3K9Me3, GSK126 treatment group showed some impact on this particular epigenetic modifier, as illustrated in Figure 5, which shows that GSK126 caused a slight increase in H3K9Me3 level. It is known that H3K9Me3, a constitutive heterochromatin specific modifier and H3K27Me3, a facultative heterochromatin specific modifier has a compensatory effect therefore it is possible that the decrease of H3K27Me3 is partially compensated by slight increase in H3K9Me3.

**Figure 5.**
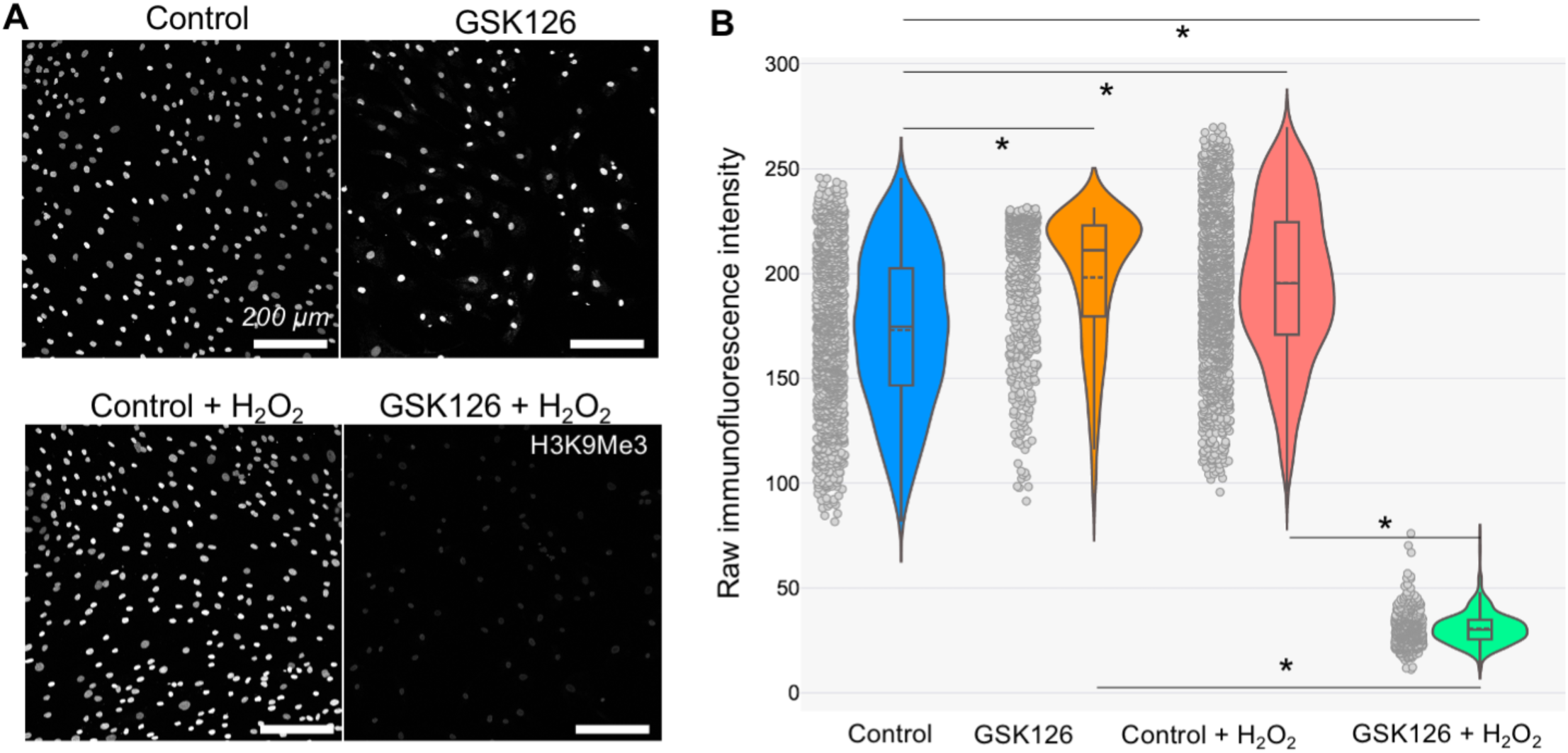
Effect of GSK126 and H_2_O_2_ treatment on the gene repression specific modifier H3K9Me3 in hBM-MSC. **(A)** Representative unprocessed image of a field of view from each group. **(B)** Data in the bar graph is based upon > 4 technical replicates (>100 cells per replicate) per group. *p <0.001 based on ANOVA.

In line with the H_2_O_2_ induced chromatin condensation and chromatin remodeling (as stated in previous sections), the H3K9Me3 intensity level increased in the control group 24 hours post-treatment with H_2_O_2_. However, in the GSK126-treated group, where chromatin remodeling was impeded, a significantly lower amount of H3K9Me3 was observed 24 hours post-treatment with H_2_O_2_, suggesting a comprehensive alteration in epigenetic status within the nucleus. It is noteworthy that in this experiment, H_2_O_2_ treatment persisted for 90 minutes, followed by complete removal of H_2_O_2_ and replenishment with fresh MSC medium. Similarly, the GSK126-treated group received fresh MSC medium containing GSK126.

### GSK126 pre-treatment changes the DNA damage profile post H_2_O_2_ treatment

Reactive Oxygen Species are known to induce DNA damage, particularly through double-strand breaks (DSBs). Cells employ various strategies for DSB repair, with an increase in γ-H2AX (phosphorylated form of H2AX) levels and foci serving as markers of this repair response. Following 24 hours of H_2_O_2_ treatment, the overall intensity level of γ-H2AX exhibited a nearly 25% decrease as a result of GSK126 pre-treatment, as shown in Figure 6A, and further quantified in Figure 6B. It is noteworthy that right after the application of H_2_O_2_, at 0 hour timepoint the DNA damage repair marker intensity level is similar in both control and GSK126 groups, and a little higher in GSK126 group.

**Figure 6.**
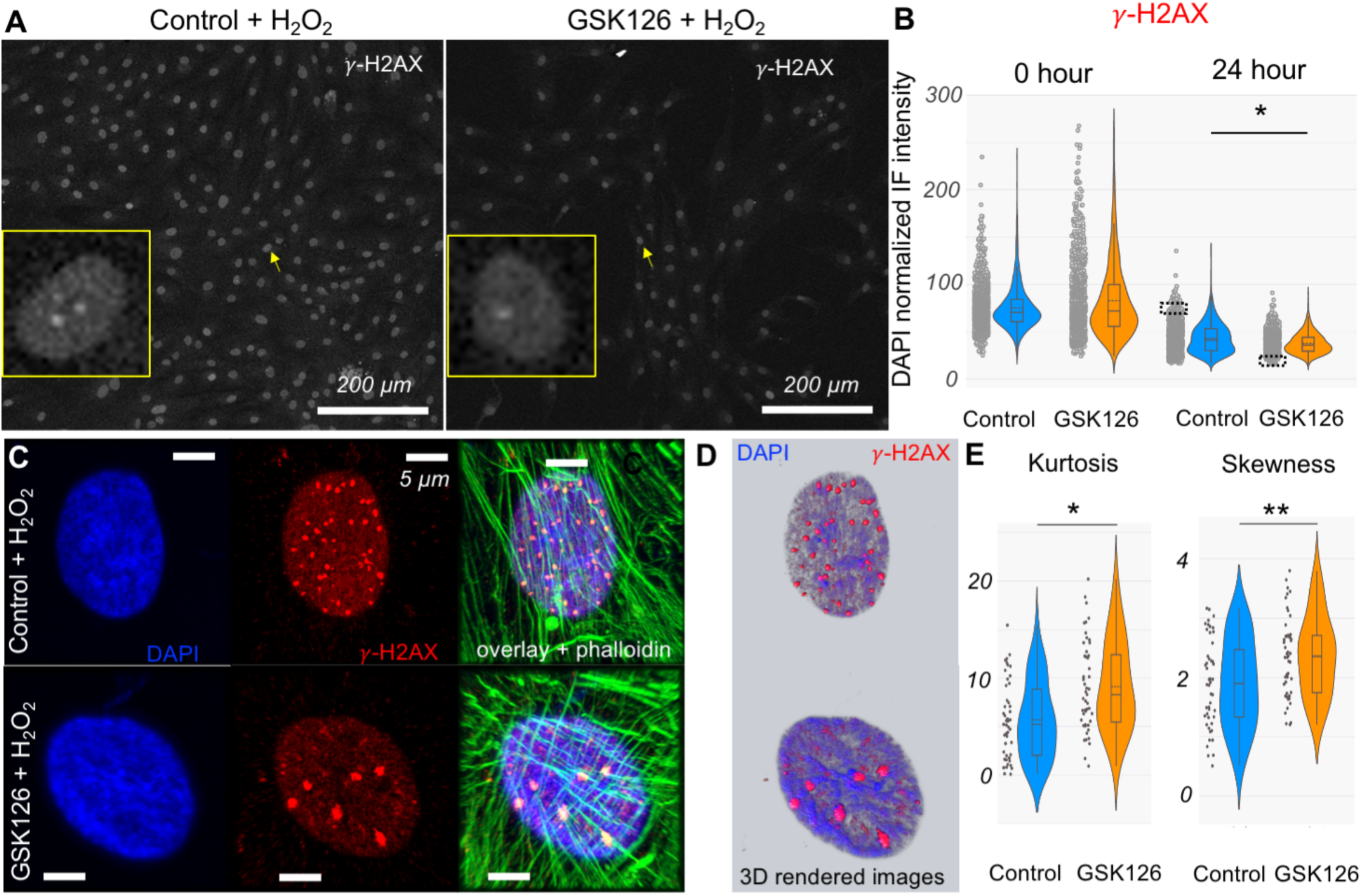
DNA damage in hBM-MSC on the exposure of H_2_O_2_. **(A)** Representative field of view of nuclei with the DNA damage marker γ-H2AX. Zoomed in view of a nucleus in each case is shown in the inset for the nucleus marked with an arrow. Dots represent the DNA damage foci. **(B)** Violin plot is based upon >4 technical replicates (>200 cells per replicate) per group. *p <0.01. **(C)** Super-resolution images of representative nucleus with multiple stains. The cases represent the extreme situations in the rightmost violin plot, and the nuclei of interest were picked from the black dotted rectangle regions. **(D)** 3D rendering of the super-resolution images based upon z stacks showing the larger foci and smaller nanoscale foci. **(E)** Kurtosis and skewness of the γ-H2AX immunofluorescence signal from individual nuclei. Each dot represents kurtosis and skewness value from one nucleus. Data based upon >45 randomly selected cells. *p< 0.001, **p = 0.002.

To understand the reason of lower intensity of γ-H2AX in GSK126 treated cells we visualized the chromatin γ-H2AX using super high-resolution microscopy. While the distribution of the number of γ-H2AX foci appeared highly heterogeneous across all fields of view, the elevated intensity of γ-H2AX immunofluorescence correlated with a greater number of γ-H2AX foci. Overall, in the control group, a greater number of cells displayed many γ-H2AX foci and a smaller number of cells displayed a smaller number of foci. In the GSK126 treated group, cells displayed the opposite trend where a greater number of cells displayed a smaller number of foci and a smaller number of cells showed plenty of foci. As illustrated in Figure 6C, a representative case from the control group exhibited numerous visible γ-H2AX foci, whereas the nucleus from the GSK126-treated group displayed fewer foci, albeit larger in size. Further analysis through three-dimensional rendering of z-stack images (145 m step size) revealed that most foci were predominantly located in euchromatin zones, which are lightly stained with DAPI, as shown in Figure 6D. Additionally, many nano-sized globules of γ-H2AX foci were visible in both groups. We quantified such intranuclear distribution of γ-H2AX by calculating the skewness and kurtosis of the γ-H2AX signal in the nucleus. Both kurtosis and skewness (Figure 6E) was higher in the GSK126 treated groups which means that intensity distribution was more clustered in spatial pockets in that group.

### Cell viability post H_2_O_2_ exposure is not affected by the GSK126 pre-treatment

Subsequently, we explored the impact of GSK126 and H_2_O_2_ on cell viability (Figure 7). Treatment with 20 µM GSK126 did not adversely affect cell viability 24 hours post H_2_O_2_ treatment, suggesting the absence of longer-term toxic effects at this specified concentration. Exposure to 500 µM H_2_O_2_ for 90 minutes did not lead to a significant decrease in cell viability after 24 hours post H_2_O_2_ treatment. Likewise, pre-treatment with GSK126 did not exert any notable influence on cell viability, even following exposure to H_2_O_2_.

**Figure 7.**
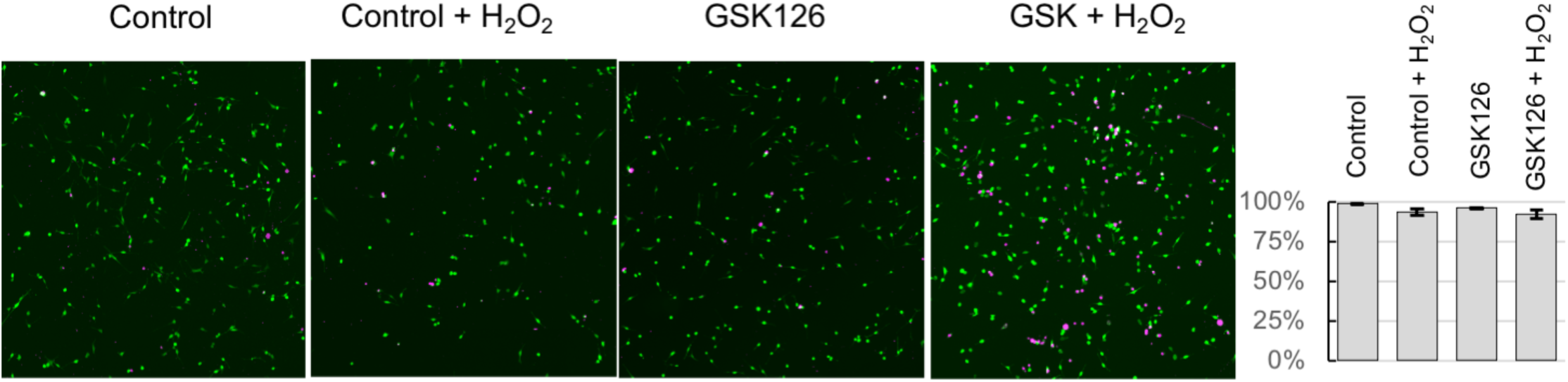
Viability of hBM-MSC on the exposure of H_2_O_2_. Representative images of the field of view from each group. Green represent all cells and pink represent dead cells. Bar graphs show percentage of cell viability in each group. Bar graphs are based upon >4 technical replicates (>1000 cells per replicate) per group.

## DISCUSSION

Overall, this study creates a technical framework to visualize and quantify the chromatin remodeling in real time, and a conceptual framework to understand the role of chromatin remodeling upon hyper-oxidative stress. Chromatin remodeling and its functional significance is a budding research niche and a few groups started working on imaging such events[36]. The technique presented in this work can be applied to many applications in other cell types where real time visualization of chromatin remodeling can reveal new fundamental discoveries. A limitation of this technique is the use of NucBlue (a live imaging formulation of Hoechst) which is a weaker stain, and it is not the optimal dye to image intranuclear architecture for a longer timeframe because of the risk of photobleaching. Endogenous probes such as fluorescent H2B coding gene can be integrated to the chromatin for long-term imaging with higher spatial resolution[37]. With that modality of imaging, more ambitious questions can be answered such as how the chromatin compaction happens in 3D. By using those probes, it is possible to capture the full 3D volumetric map of the nucleus. Additionally, in the present work the third dimension was very small because the cells are flat, cultured on a rigid polymer. Therefore, the 2D representation of the CRI mapping is valid but 3D imaging with multiple images in the z stack will provide us with more insight into the spatial distribution of CRI map.

We showed that hyperoxia results in spatially heterogeneous chromatin remodeling and compaction. Previous in situ studies with isolated DNA indicated that chromatin condenses on the exposure to oxidative stress but to our knowledge this is the first study to show such phenomena in intact cells and to quantify the remodeling at the nanoscale. Our observation of persistent and higher euchromatin remodeling is intriguing but it is not clear if this phenomenon is biophysical or biochemical, and the functional meaning of such remodeling in the euchromatin region. Interfacial phenomenon at the heterochromatin-euchromatin boundary is an emerging area of study. Further studies are required to discover any mechanism that might be underlying such observation.

EZH2 inhibition by GSK126 resulted in the loss of chromatin compaction and remodeling on the application of hyper oxidative stress. This observation might have multiple implications. One possibility is the requirement of rapid H3K27 methylation during the chromatin remodeling which is hindered by the EZH2 inhibition. Direct visualization of H3K27 methylation using a live probe may confirm such a possibility. Another possibility is that EZH2 inhibitions resulted in a higher ARID1A level which ejects and slides nucleosomes by being part of the SWI/SNF CRC thus pointing to a role of chromatin remodeling by ARID1A. Previous studies with western blot showed that in ovarian cells ARID1A expression increases by a modest amount on the application of GSK126 which matches with our observation[31]. It is not clear if ARID1A maintains the structural integrity of the chromatin or if it is actively engaged in the chromatin remodeling, or both. Therefore, it is possible that ARID1A activity becomes overtly higher because of GSK126 treatment which keeps the chromatin open. Direct inhibition of ARID1A and other structural units of the SWI/SNF CRC by RNA interference may further elucidate the process.

We observed several epigenetic modifications associated with the GSK126 treatment. The decrease in H3K27Me3 is not surprising given that it is facultative heterochromatin, and the methylation is performed by PRC2, which is catalyzed by EZH2. The slight increase in H3K9Me3 level by GSK126 can be explained by the fact that the H3K9Me3 is a constitutive heterochromatin mark, and it can compensate the loss of H3K27Me3. The drastic decrease in H3K9Me3 post hydrogen peroxide treatment for GSK126 pretreated group is intriguing and needs further investigation. It is possible that the observed lack of chromatin remodeling and chromatin compaction in the GSK126 treated group upon hyperoxia forces the chromatin to stay open by a compensatory mechanism that may employ the decreasing of H3K9Me3. Of course, this would have consequences on the MSC transcriptome and functionality which can be revealed by several assays such as RNA seq and ChIP-seq.

DNA damage foci formation was revealed by super-resolution microscopy. Interestingly, the GSK126 treatment group showed a modest decrease in the γ-H2AX formation after the oxidative stress was applied. This might have implications with the degree of DNA damage upon H_2_O_2_ application. As stated in the introduction, the role of chromatin remodeling after DNA damage is debated and one possibility indicates that damage repair molecules need access to the damaged lesions for an efficient repair. Because γ-H2AX is a marker for the DNA being actively repaired it is possible that in the GSK126 group where the chromatin stayed open, the DNA damage was mostly repaired 24 hours post stress thus resulting in an overall lower level of γ-H2AX. It is not clear if this is detrimental or beneficial for cells because cell viability did not change in the GSK126 group after the application of hyperoxia. Repeated hyper oxidative damage might cause irreversible DNA damage and senescence. If GSK126 pretreatment can help in efficient DNA damage repair, this may have implications in lowering DNA damage and senescence, but more experiments are required to elucidate such possibilities. The fact that the DNA damage foci were larger in many nuclei of GSK126 inhibited group can be explained as follows. It is possible that without GSK126, the chromatin compacts and many DNA damage lesions form which stay apart from each other due to the heterochromatin barrier created by the chromatin compaction and remodeling. When the chromatin is more open, as in the case of GSK126 group, the damaged lesions coalesce by phase separation to create larger foci. Phase separation in chromatin has been reported recently and our observation might be in line with several other studies [38].

The present work on chromatin remodeling under hyperoxia condition can be compared with previous works on chromatin level changes[39,40] under the other extreme situation of hypoxia. Hyperoxia and hypoxia, despite being opposites in terms of oxygen availability (hyperoxia being an excess and hypoxia a deficiency), share several convergent features, particularly regarding their effects on cellular physiology, oxidative stress, and the inflammatory response. Both hyperoxia and hypoxia can increase ROS production and cause oxidative stress, leading to cellular damage and the activation of protective pathways like hypoxia-inducible factor (HIF) and nuclear factor erythroid 2–related factor 2 (Nrf2). While hypoxia itself does not directly increase ROS, the reperfusion (restoration of oxygen) following hypoxic conditions can lead to a sudden burst of ROS, known as ischemia-reperfusion injury. Cells struggling to maintain homeostasis in low oxygen also experience a reduction in antioxidant defenses. Hypoxia induces profound changes at the chromatin level, altering the epigenetic landscape in ways that affect gene expression, cellular differentiation, and oxygen sensing. Studies show that histone demethylases such as KDM6A and KDM5A act as direct sensors of oxygen, and their reduced activity under low oxygen levels leads to rapid increases in histone methylation marks like H3K27. These changes occur independently of hypoxia-inducible factors (HIFs) and involve the persistence of repressive histone modifications, such as H3K27me3, which block differentiation by preventing the necessary transcriptional reprogramming. In addition, chromatin remodeling factors like SWI/SNF and histone acetyltransferases (HATs) play crucial roles in the dynamic interplay between chromatin structure and hypoxia signaling, further fine-tuning cellular responses. These previous studies combined with the present work warrants further investigation of the specific sensor in the chromatin that responds to the high oxygen level. Because we did not find any significant change in the DNA damage level immediately after the ROS activity in GSK126 compared to control groups, it is possible that the chromatin condensation effects we found might be triggered independent of DNA damage, but such speculation can possibly be tested by live visual probes for epigenetic markers and modulating several epigenetic modifiers upstream of histone methylation which is a focus of our future work.

This study potentially opens several new questions at the crossroads of DNA damage and epigenetic modifications at several levels such as chromatin remodeling at the multi-nucleosome level and histone modifications, caused by an external perturbation, in this case, hyper-oxidative stress. However, many other perturbations might cause a similar effect, which requires further investigation. Future works will focus on the functional meaning of the drastic epigenetic change as shown by the loss of H3K9Me3 upon H_2_O_2_ treatment in GSK126 treated group. Because GSK126 pre-treatment doesn’t cause lower cell viability and results in overall lower DNA damage, this might have implications in MSC based regenerative medicine therapy if the functionality of MSC is enhanced, or at least, maintained.

## EXPERIMENTAL SECTION

### Cell culture and pharmacological treatment

Human bone marrow derived primary MSC (hBM-MSC) were used for this study according to our previously reported work[33]. Cells from healthy adults were obtained from Lonza at passage 2 (PT-2501). Cells were subcultured using Lonza MSC general medium (MSCGM) consisting of MSC basal medium and growth supplements (MCGS) and maintained at 37°C, 90% humidity and 5% CO_2_. Trypsin EDTA (Lonza) were used for passaging. For all live experiments, cells were maintained in the MSCGM. For all imaging experiments, ibidi 8 well µ-slides (ibidi 80826) treated for cell culture and containing a thin polymer coverslip at the bottom for imaging were used. Cells were seeded at 5000/ cm^2^. Cells were exposed to the drug GSK126 at 20 µM concentration (Sigma Aldrich), one day post seeding and attachment. The control groups contained DMSO to match the drug’s dilutant concentration of DMSO (VWR). After 2 days of drug treatment, cell proliferation was quantified and after 4 days of drug treatment, cells were used for the hydrogen peroxide exposure study. Hydrogen peroxide was diluted in DPBS to prepare a 500 µM solution in MSC medium, with or without GSK126 as required for the experiments. Separately, murine fibroblast cell line NIH 3T3 cells were cultured for the same experiments according to our previously published work[41].

### Quantification of cell proliferation and cell viability

After 3 days of complete cell proliferation (1-day pre drug treatment, 2 days post GSK126 addition) cells were fixed with 4% PFA (in 1X PBS: Phosphate Buffer Solution) for 7 minutes at room temperature. DAPI (Thermo Fisher) was used to stain the nuclei. Images were captured at multiple, randomly selected locations. Subsequently, the images were thresholded in ImageJ and the particle analyzer was used to count the number of cells in the fields of view. That number was used to extrapolate the cell number per well (total area per well is 1 cm^2^). Then the initial cell seeding number (5000/ cm^2^) was used to calculate the fold change of cell number at day 3 compared to the initial number of seeded cells. Cell viability for all cases was performed using a live/dead assay kit (Thermo Fisher) consisting of Calcein AM and ethidium homodimer according to the manufacturer’s protocol. Cells were imaged using a confocal microscope (Zeiss LSM 980) in the 488 nm (green) and 561 nm (red) channels. The required ratio of Calcein AM and ethidium homodimer was calibrated in a preliminary experiment for the specific cell type.

### Quantification of cell and nuclear phenotype & immunofluorescence staining

Fixed cells were stained with phalloidin and DAPI (Thermo Fisher) to quantify the cell and nuclear phenotype. Images were acquired using a Zeiss LSM 980 microscope. ImageJ was used to characterize the cell and nuclear geometric parameters such as area, perimeter, P/A, solidity and circularity as described in a previous study[42]. The fluorescence intensity of the F-actin was quantified using the “plot profile” tool. For immunofluorescence studies, cells were permeabilized with 0.1% Triton X-100 in 1X PBS, washed with 1X PBS, blocked for non-specific binding sites with bovine serum albumin and normal goat serum. Primary antibodies were diluted at the required concentration in antibody dilution buffer and incubated overnight at 4°C. Next, the secondary antibody was applied at the required concentration at room temperature for 2 hours. DAPI and phalloidin staining were then performed as required. Cells were imaged using a confocal microscope (Zeiss LSM 980) using several objective lenses (10×, 20×, 40× water objective or 63× oil objective) as required for the specific experiment. Super-resolution microscopy was performed using the Airyscan 2 module of the confocal microscope and 63× oil objective. The following antibodies were used for this study. Conjugated antibodies: H3K4Me3 (CST, 9751, 1:100), H3K27Me3 (CST, 9733, 1:100); Unconjugated primary antibodies: H3K9Me3 (Invitrogen, PA5-31910, 1:400), Phospho-Histone H2A.X (Invitrogen, MA1-2022, 1:200); Secondary antibodies: Alexa Fluor highly cross-adsorbed (Invitrogen A32731, Invitrogen A32733 and Invitrogen A32727, all in the ratio 1:200).

### Live imaging of chromatin remodeling

To image the live cell response to hydrogen peroxide, NucBlue was added to the ibidi plate wells (2 drops/ ml) and maintained inside the incubator for 20 minutes at 37°C, 90% humidity and 5% CO_2_. After, the plate was placed on the confocal microscope stage and subsequent imaging was performed using the 40× water objective lens for high resolution imaging with the 405 nm laser. The sample was maintained at 37°C, 90% humidity and 5% CO_2_ during the imaging session using an incubation chamber. After choosing a suitable nucleus, the crop area was increased to visualize a single nucleus with detailed chromatin architecture. Once the nucleus was decided, an image was captured before adding the hydrogen peroxide. At this stage, the medium was pipetted out and medium containing H_2_O_2_ was added (with or without GSK126). Immediately after, the nucleus was imaged again. Images were then captured every 5 minutes for 80 minutes.

### Quantification of chromatin remodeling

After the live imaging was performed, the images were cropped and prepared for further postprocessing using ImageJ. A custom Matlab code was developed for quantification of the chromatin remodeling index. The CRI was calculated at every timepoint with respect to the image corresponding to the pre-H_2_O_2_ treatment. The images were registered using a series of in-house Matlab function instructions. This step registered each pixel of the pre-H_2_O_2_ treatment image with the pixels of the post-H_2_O_2_ treatment timepoints. The result was a set of coordinates (*X_t_, Y_t_*) at every post-H_2_O_2_ treatment timepoint and initial coordinate (*X_i_, Y_i_*) of the pre-H_2_O_2_ treatment. Accordingly, the CRI was calculated by using the following equation:
*CRI = [(X_i_ − X_i_)^2^ + (Y_i_ − Y_i_)^2^]^1/2^*

Note that this quantity is in pixels which can be converted to microns. Before the registration code was applied, an image drift correction algorithm was implemented to the image stack using ImageJ to account for any bulk drifting of the nucleus. Therefore, the CRI only accounted for the real chromatin motion, and discarded any other movement of the bulk nucleus. This technique is limited only by image resolution. We define the displacement map as a chromatin remodeling index map for this specific context. This technique to quantify the displacement inside an image from the texture of images has been thoroughly validated in previous studies[43,44]. Segregation of chromatin domains to define the euchromatin and heterochromatin regions was performed to quantify CRI in the intranuclear space. This step was performed based on a previously published work which uses a Hill’s function[35]. The pixels of the raw image were sorted based on the grayscale pixel intensity, which can have any integer value between 0 and 255, where 0 is absolute black depicting lack of chromatin and 255 is absolute white depicting highly condensed heterochromatin. Subsequently, a user independent automated algorithm was utilized to segregate the euchromatin, interchromatin and heterochromatin domains.

### Statistical Analysis

All data is reported for all experiments as raw data and no outlier was removed. Two-sided student’s t-test was used to compare two groups. ANOVA was used to quantify differences between multiple groups follows by post-hoc Tukey’s test, as appropriate. Normality of the dataset was confirmed first before applying the t-tests. Error bars in bar graphs represent mean ± standard deviation. The sample sizes which include the number of technical replicates, number of cells (when applicable), and p-values are reported in each individual figure. For all statistical analysis R was used.

## Acknowledgements

The authors acknowledge the funding from the National Science Foundation (CAREER award, 2236710), which partially supported this work. We acknowledge Abigail Fennell, Zack Aboellail, Jack Forman and Lillian Garfinkel for their contribution in imaging and data analysis, and Scott Burlingham for insightful discussions.

## Author contributions

S.G. performed conception and design. L.M., N.K., N.C. and S.G. performed experiments and data collection. L.M., N.K., S.K., N.C. and S.G. performed data analysis and interpretation. L.M. and S.G. performed drafting and revising of manuscript. S.G. performed resources and funding acquisition.

## Data Availability Statement

The data that support the findings of this study are available from the corresponding author upon reasonable request. The code to quantify chromatin remodeling can be found here: https://github.com/GhoshLabEng/Chromatin-Remodeling

