## Supplementary File for "Hyperoxia Induced Alteration of Chromatin Structure in Bone Marrow Derived Primary Mesenchymal Stromal Cells"

Lauren A. Monroe, Samantha Kaonis Natalie Calahan, Neda Kabi, Soham Ghosh
Translational Medicine Institute
Colorado State University
2350 Gillette Dr, Fort Collins, CO 80523, USA

Natalie Calahan
Department of Chemical and Biological Engineering
Colorado State University
400 Isotope Dr, Fort Collins, CO 80521, USA

Soham Ghosh
Department of Mechanical Engineering
Colorado State University
400 Isotope Dr, Fort Collins, CO 80521, USA

Soham Ghosh
Cell and Molecular Biology
Colorado State University
1050 Libbie Coy Way, Fort Collins, CO 80524, USA

### SUPPORTING INFORMATION

#### Supplementary Figures

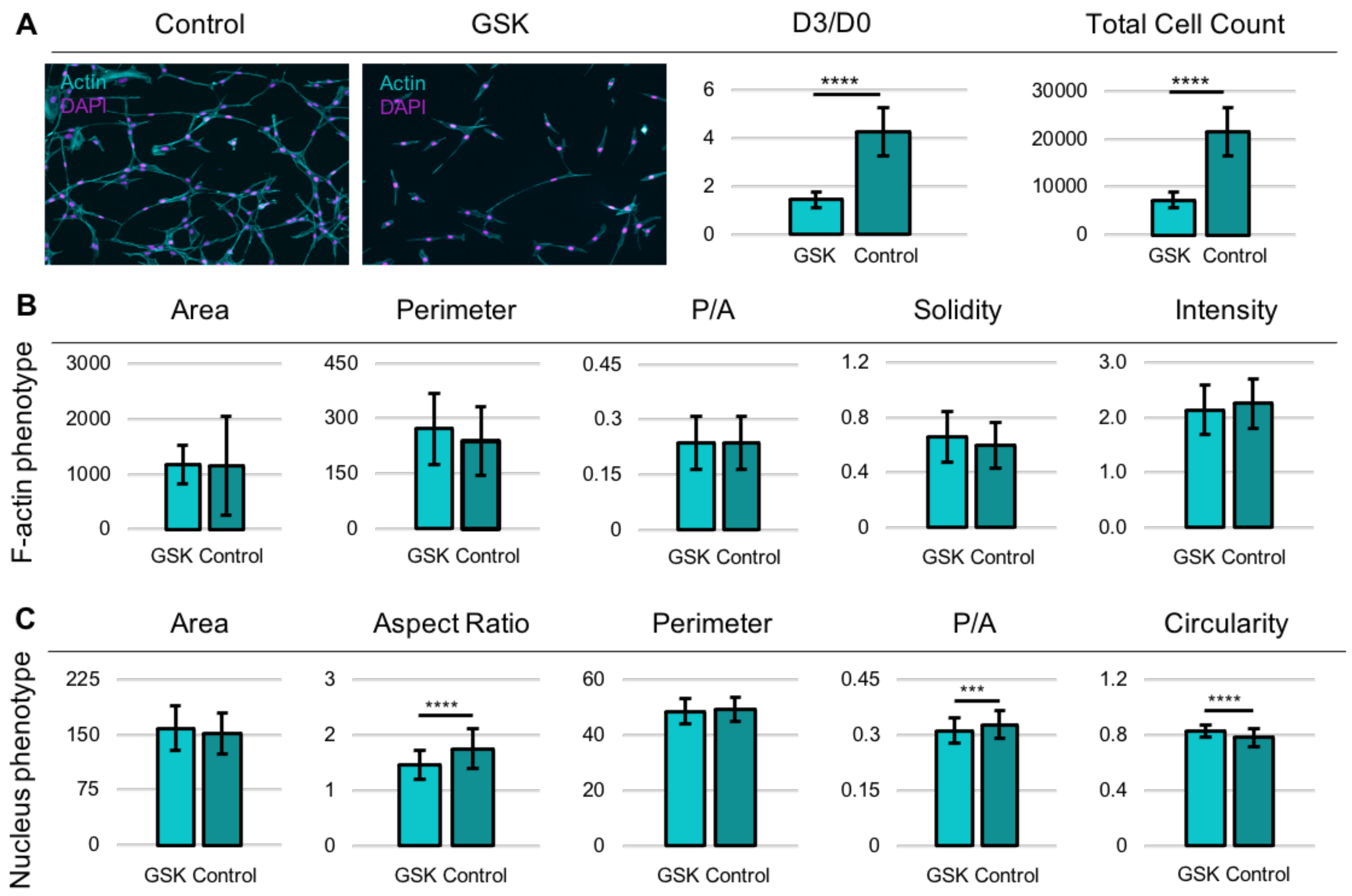


**Figure S1. Effect of 20 µM GSK126 treatment on the proliferation and phenotype of NIH 3T3 murine embryonic fibroblasts. (A)** Representative images of Passage 4 MSC with phalloidin stained F-actin and DAPI stained nucleus on Day 3 after seeding. Ratio of number of cells on Day 3 over Day 0. Total number of cells on Day 3. **(B)** Different geometric shape parameters based upon cell phenotype. **(C)** Different geometric shape parameters based upon nuclear phenotype. Data based upon >4 technical replicates (>2000 cells per replicate) per group. ****p <0.0001, ***p <0.001.

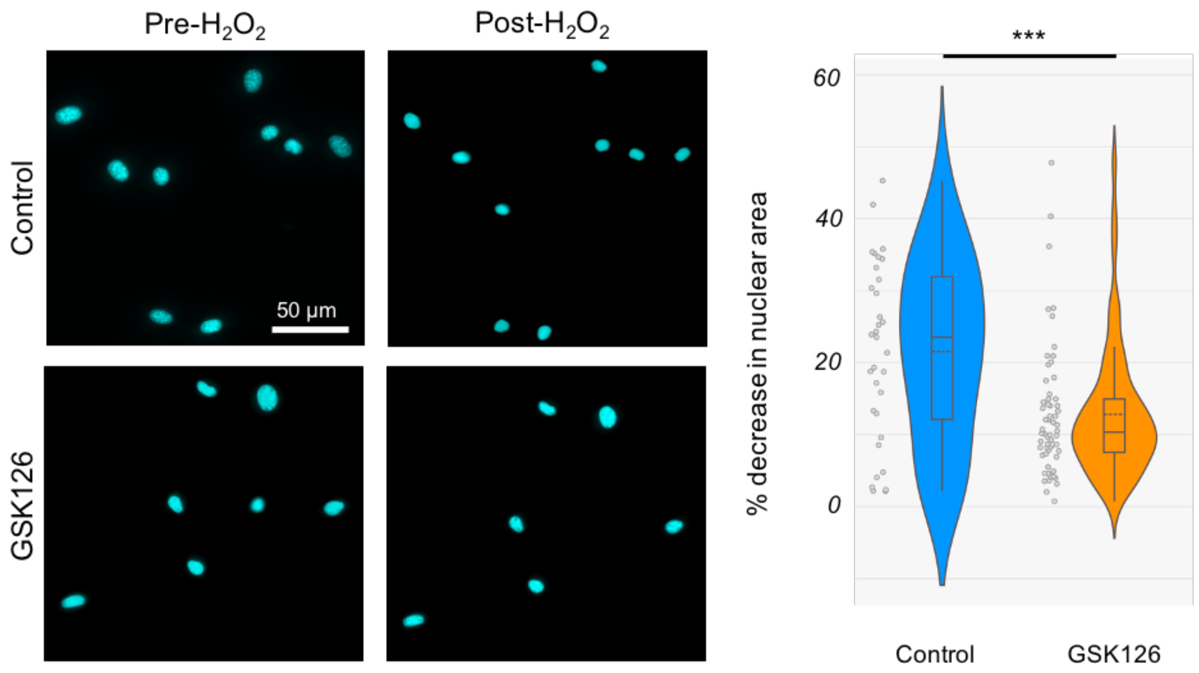


**Figure S2. $H_2O_2$ (500 µM) driven nuclear shrinkage in NIH 3T3 cells.** Representative Images of Nucblue stained nuclei in live cells. Post-$H_2O_2$ corresponds to 75 min timepoint. Data based upon > 50 cells per group. *p <0.01.
